# Sex-dependent dominance at a single locus maintains variation in age at maturity in Atlantic salmon

**DOI:** 10.1101/024695

**Authors:** Nicola J. Barson, Tutku Aykanat, Kjetil Hindar, Matthew Baranski, Geir H. Bolstad, Peder Fiske, Céleste Jacq, Arne J. Jensen, Susan E. Johnston, Sten Karlsoon, Matthew Kent, Eero Niemelä, Torfinn Nome, Tor F. Næsje, Panu Orell, Atso Romakkaniemi, Harald Sægrov, Kurt Urdal, Jaakko Erkinaro, Sigbjørn Lien, Craig R Primmer

**Affiliations:** Centre for Integrative Genetics (CIGENE) and Department of Animal and Aquacultural Sciences, Norwegian University of Life Sciences, NO-1432 Ås, NORWAY; Department of Biology, University of Turku, FI-20014, FINLAND; Norwegian Institute for Nature Research (NINA), NO-7485 Trondheim, NORWAY; Nofima AS, PO Box 210, NO-1431 Ås, NORWAY; Natural Resources Institute Finland, Oulu FI- 90014, FINLAND; Institute of Evolutionary Biology, University of Edinburgh, Charlotte Auerbach Road, Edinburgh, EH9 3FL, UNITED KINGDOM; Radgivende Biologer, NO-5003 Bergen, NORWAY

## Abstract

Males and females share many traits that have a common genetic basis, however selection on these traits often differs between the sexes leading to sexual conflict^1,2^. Under such sexual antagonism, theory predicts the evolution of genetic architectures that resolve this sexual conflict^2-6^. Yet, despite intense theoretical and empirical interest, the specific genetic loci behind sexually antagonistic phenotypes have rarely been identified, limiting our understanding of how sexual conflict impacts genome evolution^4,7,8^ and the maintenance of genetic diversity^8,9^. Here, we identify a large effect locus controlling age at maturity in 57 salmon populations, an important fitness trait in which selection favours earlier maturation in males than females^10^, and show it is a clear example of sex dependent dominance reducing intralocus sexual conflict and maintaining adaptive variation in wild populations. Using high density SNP data and whole genome re-sequencing, we found that vestigial-like family member 3 (*VGLL3)* exhibits sex-dependent dominance in salmon, promoting earlier and later maturation in males and females, respectively. *VGLL3*, an adiposity regulator associated with size and age at maturity in humans, explained 39.4% of phenotypic variation, an unexpectedly high effect size for what is usually considered a highly polygenic trait. Such large effects are predicted under balancing selection from either sexually antagonistic or spatially varying selection^11-13^. Our results provide the first empirical example of dominance reversal permitting greater optimisation of phenotypes within each sex, contributing to the resolution of sexual conflict in a major and widespread evolutionary trade-off between age and size at maturity. They also provide key empirical evidence for how variation in reproductive strategies can be maintained over large geographical scales. We further anticipate these findings will have a substantial impact on population management in a range of harvested species where trends towards earlier maturation have been observed.

---

The importance of balancing selection in maintaining variation in fitness related traits, which are expected to be under strong selection, is a long standing question in evolutionary biology^9,14,15^, with recent models suggesting that balancing selection may be particularly important in maintaining genetic variation^11,16^. Sexually antagonistic selection on traits with a shared genetic architecture where each sex is displaced from their phenotype optima is one mechanism generating balancing selection^2,4,8,11,17^. Theoretical models predict that dominance reversals, where the dominant allele in one sex is recessive in the other, greatly reduce constraints on the resolution of sexual conflict and may be particularly efficient at maintaining variation, as it can result in heterozygote superiority across the sexes^8,15,18^ but this has never been observed in the wild. A paucity of empirical examples with known genetic architecture means that the evolutionary significance of sexual conflict, and its subsequent resolution or persistence, remains largely unknown^3,17,19,20^. Determining the genetic architecture of phenotypes under sexually antagonistic selection would be a key advance in our understanding of its importance in maintaining adaptive variation^4,8,19,20^.

The age at which an individual reproduces is a critical point in its life history. Age at maturity affects fitness traits including survival, size at maturity and lifetime reproductive success^21,22^. Age at maturity in Atlantic salmon (*Salmo salar*) represents a classic evolutionary trade-off: larger, later maturing individuals have higher reproductive success on spawning grounds^23,24^, yet also have a higher risk of dying prior to first reproduction^21^. Atlantic salmon reproduce in freshwater, with offspring migrating to the sea to feed before returning to their natal river to spawn. The number of years spent at sea before spawning, i.e. their age at maturity, or ‘sea age’, has a dramatic impact on size at maturity, typically 1-3 kg and 50-65 cm after one year compared to 10-20 kg and >100 cm after three or more years^25^. Larger size, and therefore later maturation, correlates closely with fecundity in females in particular and there is evidence for sex-specific selection patterns on age at maturity, as life-history strategies differ considerably between males and females, both within and among populations^23,24^. On average, males mature earlier and at smaller size, whereas females mature later, with a stronger correlation between body size and reproductive success compared to males^23,24^.

We investigated the genetic basis of age at maturity in Atlantic salmon using three independent datasets. The first, *TAN*, included two intensively sampled subpopulations from a large river system (Tana/Teno River; 68-70^°^N: *n* = 463); the second, *NOR*, comprised 54 populations spanning the Norwegian coast from 59^°^N to 71^°^N, containing both Atlantic and Barents/White Sea phylogeographic lineages^26^ (*n* = 941; *NOR* mean *n* per population = 17.4). Both datasets sampled geographically proximate populations with contrasting age at maturity (Fig. 1, Supplementary Note, Extended Data Table 1). Genome-wide association studies (GWAS) for age at maturity were then conducted within both datasets using data from 209,070 polymorphic SNPs (Supplementary Note). An approximately 100 kb region on chromosome 25 was strongly associated with age at maturity in both datasets (GWAS; *P* < 1 × 10^-20^, Fig. 1b and d, Extended Data Fig. 1) explaining 32.9 % (s.d. 4.2) of total phenotypic variation (see online methods). This association was further validated in the phylogeographically distant *BAL* dataset (*P* < 9.74 × 10^-8^, Extended Data Fig. 2), thus confirming that the region is evolutionary conserved across all European lineages. The region included two candidate loci (Fig. 1d, Extended Data Fig. 3a), vestigial-like family member 3 (*VGLL3*) and A-kinase anchor protein 11 (*AKAP11*). *VGLL3* is a transcription cofactor with a role in adipogenesis as a negative regulator of terminal adipocyte differentiation, and its expression is correlated with body weight, mesenteric and gonadal adipose content in mice^27^. *VGLL3* has also been associated with age at menarche^28,29^ and pubertal height growth in humans^30^ indicating a remarkably high level of functional conservation at this locus. Age at menarche is associated with adiposity in humans^29,30^ and puberty in fish is linked to the absolute level or rate of accumulation of lipid reserves^31^. Threshold levels of fat reserves at critical times of year are thought to control the decision whether or not to delay maturation in salmon^32,33^. Therefore, *VGLL3* may serve as a mechanism to regulate the interaction between fat reserves (adiposity) and maturation in salmon, in a similar manner as in mammals and is a strong candidate gene for age at maturity. *AKAP11* is expressed throughout spermatogenesis and is important for mature sperm motility^34^.

**Figure 1.**
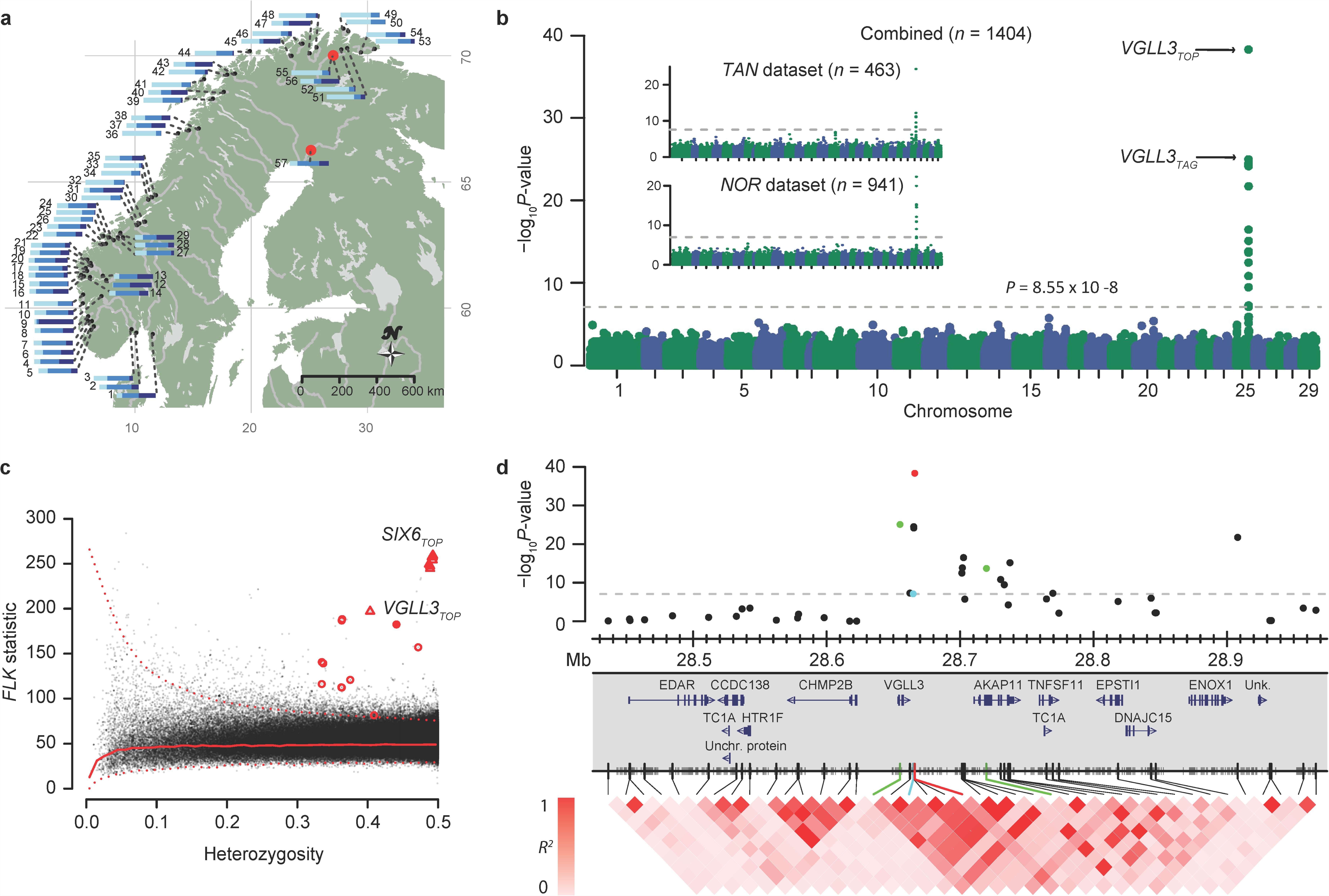
Study population details, genetic mapping of age at maturity, and divergence across populations. **a,** Map of study populations. Numbers are population IDs; 1-54: NOR, 55-56: TAN; 57: BAL datasets (see Extended Data Table 1 for details). Bars indicate the proportion of individuals maturing after one (light blue), two (medium blue) or ≥ three years (dark blue). **b,** Manhattan plots of the GWAS for age at maturity in Atlantic salmon for two independent sets combined (TAN and NOR). Inset shows the two datasets independently (see Extended Data Figure 1 for details). **c,** Signatures of spatially divergent selection using the FLK *F*_ST_ outlier test. Solid and dashed lines indicate the smoothed median and 99.5% quantile of the null (neutral) distribution. Ten SNPs flanking the *VGLL3*_*TOP*_ and *SIX6*_*TOP*_ SNPs (solid symbols) are marked with red circles and triangles, respectively. (*P*_*VGLL3TOP*_ = 1.44 × 10^-15^ and *P*_*SIX6TOP*_ ~ 0). **d** The gene model and LD plot of ~0.5 Mb region around the significant region on chromosome 25. Notable SNPs are colour coded with red (*VGLL3*_*TOP*_), blue (*VGLL3*_*iHS*_), and green (SNPs tagging missense mutations in *VGLL3* and *AKAP11*). Shorter tick marks in the SNP axis indicate resequencing variants.

A second genomic region, which spans 250 kb on chromosome 9, was strongly associated with age at maturity (*P* < 10^-20^, Extended Data Fig. 1a, Extended Data Fig. 3b-c, Supplementary information), but was no longer significant following population stratification correction (Extended Data Fig. 1). This locus is likely to represent among population variation in a correlated trait, size at maturity (See supplementary notes, Extended Data Fig. 1c-e). The core haploblock included a transcription factor of the Hypothalamus-Pituitary-Gonadal axis, *SIX6*, associated with size and age at maturity in humans^29^ and a conserved non-coding element that aligns to a candidate distal forebrain enhancer of *SIX6*^35^ (Extended Data Fig. 3b-c, Supplementary information). Both genome regions also exhibited strong signals of spatial divergent selection across populations (*FLK* outlier test, *P* < 10^-15^, Fig. 1c).

Our data were consistent with the two alleles at the *VGLL3* locus underlying age at maturity, conferring either early (*E*) or late maturation (*L*). *LL* individuals had significantly higher odds ratios for delaying maturation, particularly for older maturity ages (Fig. 2a) and were predicted to mature, on average, 0.87 (females) and 0.86 (males) years later than *EE* individuals (Fig. 3a); a remarkable shift considering the average age at maturity in salmon is 1.6 years (population range averages: 1.0 to 2.6)^25^. This locus also influenced size of individuals with the same age at maturity e.g. length = 100 and 80 cm, for *LL* and *EE* males maturing after three years at sea, respectively (*P* = 0.006, Fig. 3b, Supp. Table 1). In addition, there were striking differences in dominance patterns between the sexes: in females the late-maturation allele (*L*) was significantly partially dominant (δ = 0.38 ± 0.10, *P* = 0.033), whereas in males the early maturation allele (*E*) exhibited a strong complete dominance pattern (δ = -0.99 ± 0.16, *P* < 0.001, Fig. 2b, 3a, Extended Data Fig. 4, Extended Data Table 2), providing a compelling mechanism to explain the larger proportion of males that exhibit an early maturing phenotype compared to females^23,24^.

**Figure 2.**
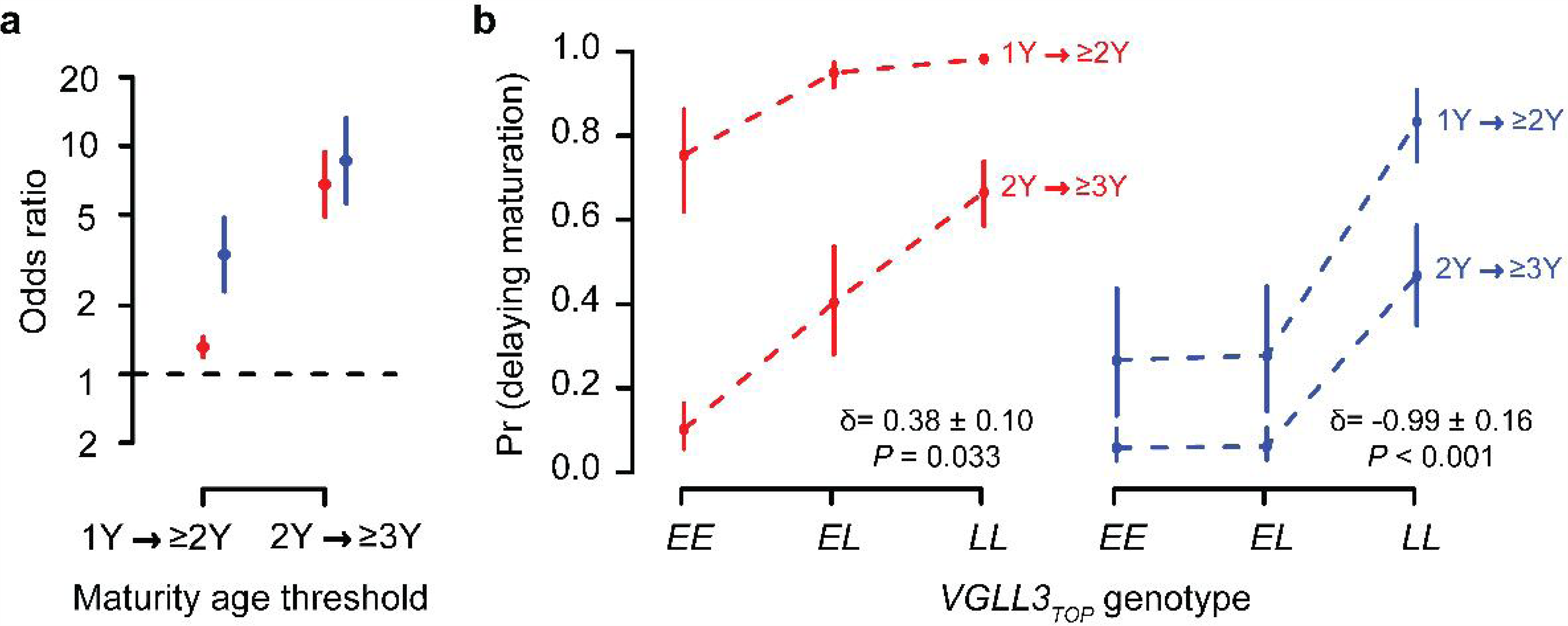
Genetic architecture of age at maturity in the *VGLL3*_*TOP*_ locus. **a,** Odds ratio (median) between the alternative homozygous genotypes for delaying maturation females (red) and males (blue). Error bars are 50% sampling quantiles (100,000 parametric permutations). All odds are significantly different from 1 (*P* < 0.001). **b,** Probability of delaying maturation as a function of *VGLL3*_*TOP*_ genotype in females (red) and males (blue). Dominance estimates in the liability scale are given for each sex (See also Extended Data Fig. 5).

**Figure 3.**
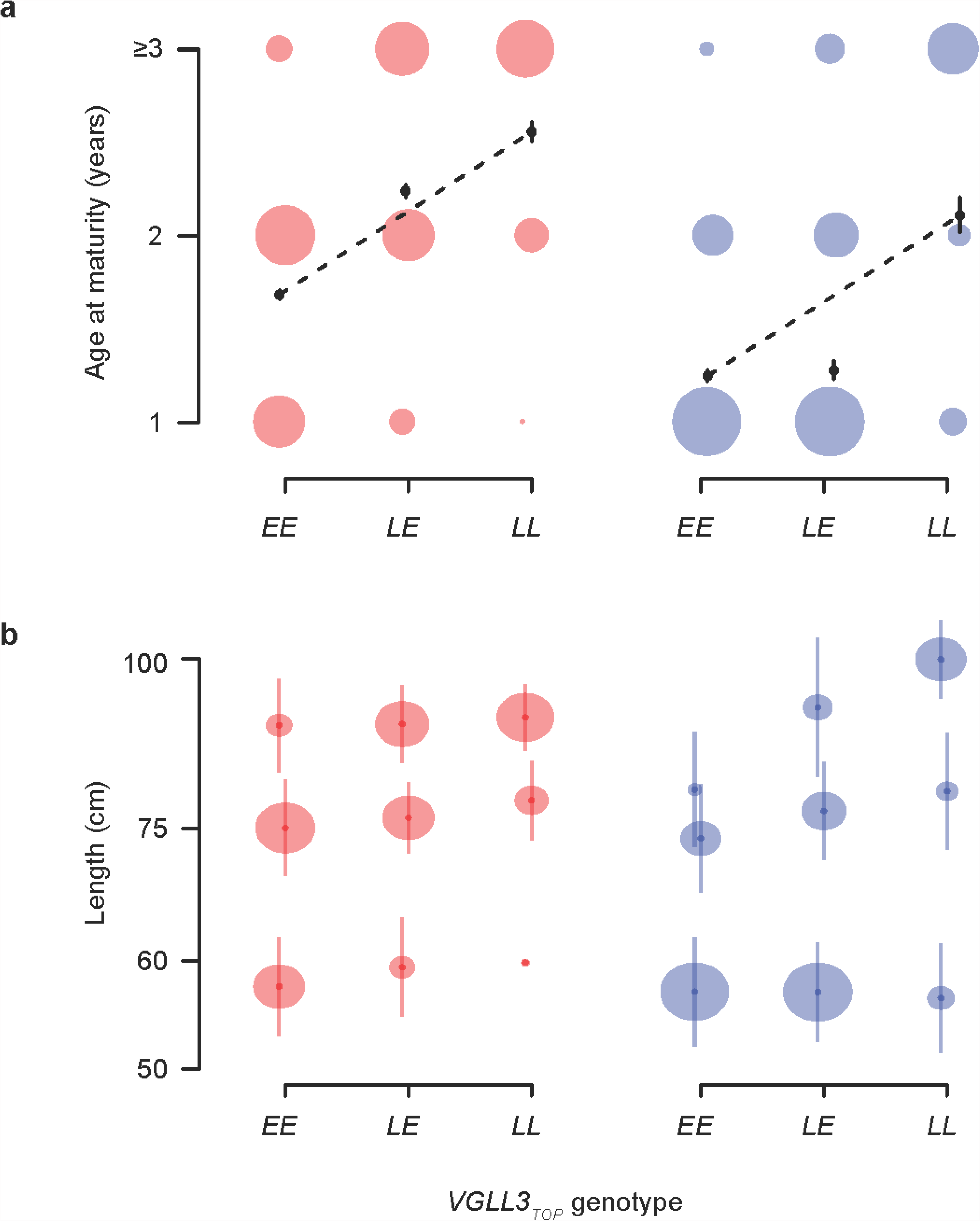
Effect of the *VGLL3*_*TOP*_ genotype on age at maturity and size. **a,** Age at maturity (number of years at sea prior to spawning migration) of females (*n* = 693, red) and males (*n* = 711, blue) in relation to *VGLL3*_*TOP*_ genotype. The circle area is proportional to sample size. Black dots indicate predicted average sea age using logit transformation model, and error bars are 50% sampling quantiles (10,000 parametric permutations). **b,** *VGLL3*_*TOP*_ genotypic effect on size maturation age classes. The average length (cm) of females (red) and males (blue) maturing after 1, 2, or 3 years feeding at sea are indicated by the lower, middle and upper three dots, respectively, in each panel. Circle diameters are proportional to sample size, and lines indicate sample standard deviation. Length (cm) on the y axis is log scaled and corrected for population effects.

**Figure 4.**
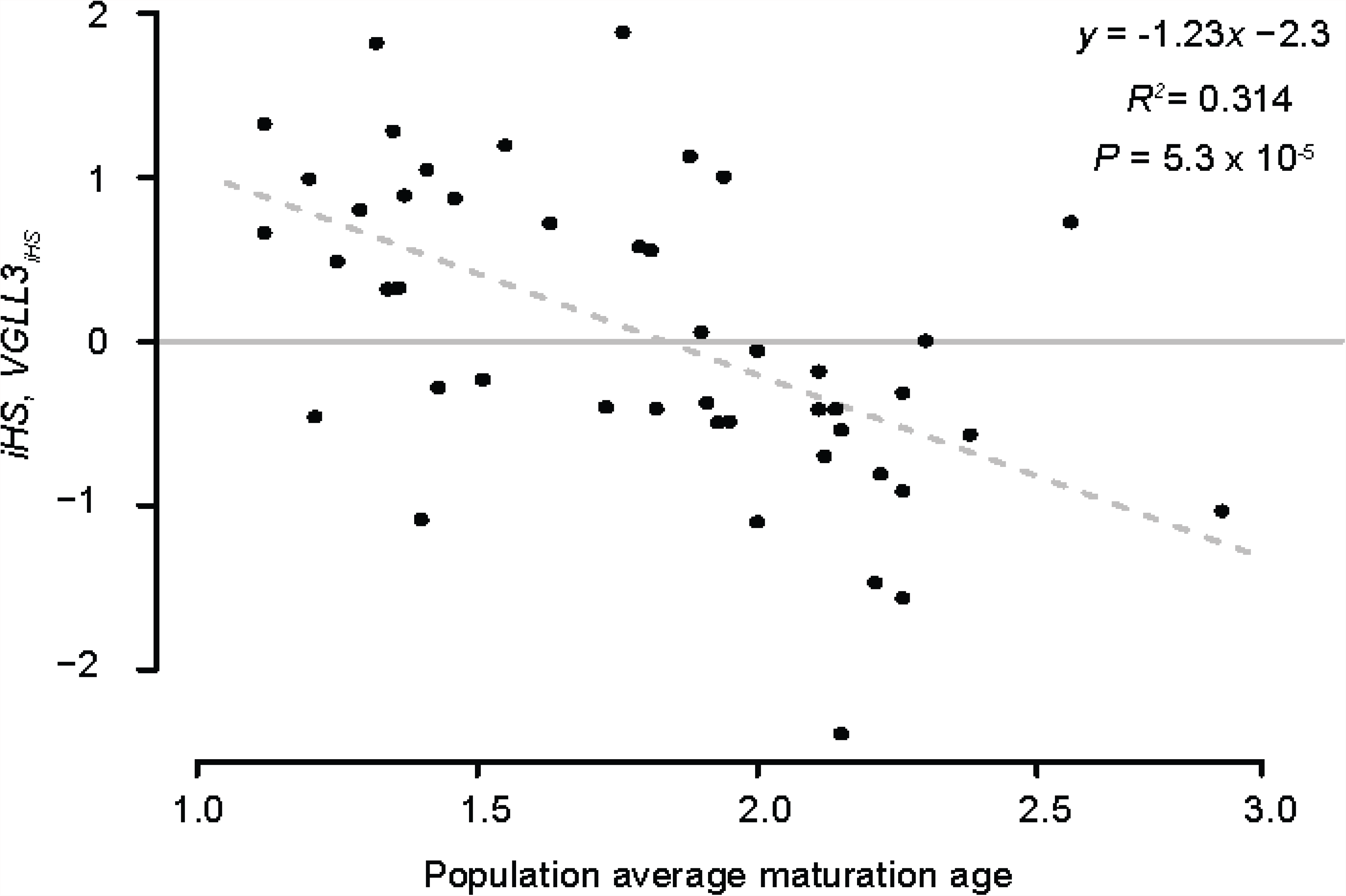
Relationship between population *iHS* score and average maturation age of each population for the *VGLL3*_*iHS*_ locus. *iHS* = 0 (no haplotype length difference) is marked with a horizontal grey line.

The mutational effect size of sexually antagonistic loci is expected to be higher when the orientation of mutational effects in the sexes is strongly decoupled ^11^, providing theoretical support for the large effect size we observed for this locus. Targeted whole genome resequencing of 32 individuals from seven populations revealed two missense mutations in *VGLL3* tightly linked to the associated SNPs (*VGLL3*_*TAG*_, 306 and 2,356 bp downstream, *R*^2^ = 1 and 0.71, respectively, Extended Data Fig. 3a) and confirmed a missense SNP had been genotyped at *AKAP11* (Fig. 1d, Extended Data Fig. 3a). A test for predicting changes in protein structure/function, PolyPhen-2 ^36^, strongly supported two of these mutations having an effect on phenotype (*VGLL3* Asn323Lys, Naïve Bayes posterior probability = 0.976, sensitivity = 0.76, specificity = 0.96; *AKAP11* Val214Met, Naïve Bayes posterior probability = 0.716, sensitivity = 0.86, specificity = 0.92). Variation at this locus was maintained in all but one of the 54 *NOR* populations, with populations characterised by large salmon consistently maintaining intermediate allele frequencies, consistent with balancing selection ^9^ (Extended Data Table 1). Given that a large proportion of variation in age at maturity (and subsequent body size) is governed by a single locus of large effect, such sex-specific trade-offs with a shared genetic basis could effectively maintain genetic variation under varying patterns of dominance between the sexes ^8^. Evolution towards complex traits controlled by fewer loci with larger effects, is also predicted where gene flow between environments with different trait optima results in balancing selection ^9,12,13,37^. Local adaptation with gene flow implies ongoing, or very recent, spatially varying selection is operating and for trade-offs at the phenotypic level to be accompanied by trade-offs at the individual locus level, otherwise a single high fitness allele would invade all environments ^12^. We investigated patterns of positive selection on the *VGLL3* locus and found a strong effect of a population’s average age at maturity on the integrated haplotype score (*iHS*) ^38^, a measure of the amount of extended haplotype homozygosity at a SNP relative to the alternative allele (slope = -1.23 ± 0.29 s.e. year^-1^, *R*^2^ = 0.314, *P* = 5.3 × 10^-5^; Fig. 4, Extended Data Fig. 5). Extended homozygosity around the *L* allele was observed in populations with an older average age at maturity, while extended homozygosity around the *E* allele occurred in populations with a younger average age at maturity (Fig. 4, Extended Data Fig. 5). This result suggests a systematic shift in positive selection for earlier/later maturation alleles coincident with the population’s average sea age and is consistent with divergent selection among populations towards local optima, and an effect of gene flow on the observed genetic architecture ^13^.

Our results reveal a major effect locus determining age at maturity in Atlantic salmon. The large effect of this locus is remarkable given that age at maturity is considered to be a classic polygenic trait in many species e.g.^28,29^. The finding of a shared gene controlling age at maturity between mammals and a teleost fish provides an example where the genes controlling life history traits show a high level of conservation across large taxonomic distances, as seen for morphological characters ^39,40^, but with very different effect sizes. Our results provide the first empirical example of dominance reversal permitting greater optimisation of phenotypes within each sex. Partial dominance of the higher fitness allele in each sex can result in a net effect of heterozygote superiority across the sexes, and thus maintain stable polymorphisms^8^. In common with many other species, Atlantic salmon lack heteromorphic sex chromosomes^43^, which precludes the use of the X chromosome to protect sexual conflict polymorphisms. Sex dependent dominance removes the restrictive conditions on maintaining conflict loci on autosomes, making sexually antagonistic polymorphism more likely to be maintained on autosomes than on the X chromosome^4,8^. In line with our results, restrictive conditions on the maintenance of variation by balancing selection suggest fewer, large effect loci will control traits under both sexual antagonism and spatially varying selection ^9,11-13,37^. The discovery of a major locus affecting age at maturity will also have a substantial impact on population management of Atlantic salmon, where a decrease in the frequency of late maturation has been observed in many populations^41^ and potentially other exploited species showing comparable shifts towards earlier maturation^42^ if this architecture is similar in other species.

## Author contributions

C.R.P., S.L., N.J.B., T.A. and K.H. conceived the study. C.R.P., S.L., N.J.B., T.A., K.H., C.J., S.K. and S.E.J. designed the experiments. M.K. and T.N., generated, and conducted bioinformatics on, the molecular data. K.H., P.F., A.J.J., T.F.N., H.S., K.U., J.E., P.O. A.R. and E.N. co-ordinated the collection of phenotypic data. T.A., N.J.B., M.B., G.H.B., S.K. and C.J. analysed the data. N.J.B., T.A. and C.R.P. wrote the manuscript. All authors read and commented on the manuscript.

## Acknowledgements

We thank Leif Andersson and Thomas Hansen for commenting on earlier drafts of the manuscript. We also acknowledge the help and cooperation of the numerous fishermen and women who contributed scales and phenotypic information. Scale age measurements were carried out by Jari Haantie and Jorma Kuusela, Irmeli Torvi, Gunnel Østborg and Jan Gunnar Jensås. We thank Silje Karoliussen for genotyping of all samples with the affymetrix SNP array, Kristina Vagonyte-Hallan for DNA sexing of NOR samples, Terese Andersstuen, Katja Salminen, Meri Lindqvist, Karin Sõstar and Terhi Pajula, and Torveig Balstad, Line Birkeland Eriksen and Merethe Spets for laboratory assistance, Mikko Ellmen, Olavi Guttorm, Topi Pöyhönen, Timo Kanniainen and Arto Koskinen for sampling assistance and Teshome Mulugeta for informatics support. Bioinformatic analyses were performed using resources at the Finnish Centre for Scientific Computing (CSC), the Abel Cluster, owned by the University of Oslo and the Norwegian metacenter for High Performance Computing (NOTUR), and operated by the Department for Research Computing at USIT, the University of Oslo IT-department, and the Orion Computing Cluster at CIGENE. This study was funded by the Finnish Academy (grants 137710, 141231, 272836, 284941) and by the Research Council of Norway (QuantEscape, grant 216105) and RCN-project; 221734/O30.

## METHODS

### Study design and study material

*Norway data-set (NOR):* Individuals for the *NOR* dataset were sampled from populations spanning the Norwegian coast from the Skagerrak in the south (59 °N) to the Barents Sea in the north (71 °N). In total 54 populations were sampled including 12 populations within the Barents/White Sea phylogeographic group^43^, the remaining belonging to Atlantic phylogeographic group. (Extended Data Table 1). The *NOR* data set was initially filtered to remove any individuals with possible aquaculture ancestry^44^.

*Tana dataset (TAN):* Two sub-populations, occurring in sympatry in the mainstem Tana River (*TAN*) were subject to in depth within population sampling (n=463). The large Tana River in Northern Europe (68–70°N, 25–27°E) supports one of the world’s largest wild Atlantic salmon populations, with up to 50,000 individuals being harvested by local fishers and recreational fisheries annually^45^. Scales were collected from salmon harvested by local fishermen using a variety of methods (nets, rod) from 2001 to 2003. In total, 326 and 137 individuals from sub-populations 1 and 2 were included^69^.

*Baltic Sea data-set:* The *BAL* dataset included 114 individuals from the Tornio river (*BAL*: (66– 69°N, 19–25°E) Fig. 1, Extended Data Table 1). This population belongs to the phylogeographically distinct Baltic salmon lineage^43^. Scales were collected from individuals harvested by trained anglers from 2003 to 2005. Storage and phenotypic measurements were as described for the *TAN* samples.

In total, the study included 1,518 Atlantic salmon individuals after initial data filtering that removed low quality samples and individuals with signs of aquaculture ancestry (Extended Data Table 1).

### Phenotypic measurements

Length at capture (*LEN*), i.e. length of the fish from the tip of the snout to the natural tip of the tail (tip of the longer lobe of the caudal fin) and weight at capture (*WGT*) were recorded during sampling. The sex (*SEX*) of most individuals was determined genetically in the *NOR* and *TAN* data-set^46^, while phenotypic sex determination was used for a small subset of samples (15% and 0.3% for *TAN* and NOR datasets respectively) and for all *TOR* individuals.

Similar to tree rings, scale growth in fishes is commonly used to infer individual growth and age (e.g.^47,48^. Here, we used scales to infer growth, freshwater age (i.e. age prior to sea migration, *FW Age*), and years spent at sea prior to first sexual maturation and spawning, referred to here as age at maturity (*Mat Age*) using internationally agreed guidelines for Atlantic salmon scale reading^49^. Early life history growth traits and size were assessed for their influence on age at maturity as it has been shown that freshwater growth may be negatively correlated with seawater growth^50^ and that freshwater size may be positively correlated with age at maturity^51^. Freshwater size (*FWS*), freshwater growth (*FWG*), as well as first year growth at sea (*SWG*) were derived from the scale data, and used as independent variables throughout the analyses. *FWS* is the log radius of the scale from the scale centre to the end of the freshwater growth period, and *SWG* is the log radius of the scale from the end of the freshwater growth to the end of the first winter annulus at sea. *FWG* is dependent on *FWS* and negatively on *FW Age*. The residuals of the linear regression between *FWS* and *FW Age* were further corrected for *FW Age* in order to obtain the *FWG* metric. As expected, a model where *FWS* was nested within *FW Age* explained 97 % of *FWG* (ANOVA, *P* < 10^-16^). Size at the end of first year at sea (*SWS*) was completely dependent on *FWS* and *SWG*, and therefore was not explored as an independent variable to avoid co-linearity.

### Genotyping and data filtering

*SNP array details:* A custom 220K Affymetrix Axiom array was used to genotype samples (*n* = 2,268) according to manufacturer’s instructions using a GeneTitan genotyping platform (Affymetrix, USA). The SNPs on this array were a subset of those included on the 930K XHD *Ssal* array developed by Moen et al., (in prep), and were chosen for maximum informativeness on the basis of their SNPolisher performance (SNPolisher, V1.4, Affymetrix), minor allele frequency (*maf*) in aquaculture samples (*maf* > 0.05), and physical distribution. The ascertainment bias of this array for wild Norwegian salmon is expected to be low because of the recent founding of the aquaculture population from a large number of (*n* = 40) Norwegian salmon populations^52^.

### Genotyping protocol and data filtering

Raw genotyping data were analyzed using the linux based APT pipeline applying best practices thresholds (DQC threshold, 0.82; STEP1, 0.97). After the initial sample filtering, markers with low *maf* (*<* 0.01) and/or call rate (< 0.97) were filtered out using the *check.marker* function in the GENABEL package (v1.8.8,^53^ in the R environment (v 3.1.0,^54^ for the GWAS. The same data parameters were used for the dataset for phasing except that no *maf* threshold was set (see below). We did not perform a Hardy-Weinberg equilibrium test, since the data set contained individuals from multiple populations. After the filtering steps, 209,454 SNPs in 29 linkage groups remained in the analysis. An additional filtering was performed separately for the *BAL* dataset (*maf* < 0.01) resulting in 167,410 SNPs remaining in this dataset for the GWA analysis.

### Model selection and genome-wide association study (GWAS) of maturation age

We performed a GWAS of age at maturity (*Mat Age*) using an additive cumulative proportional odds model with the R package ORDINAL^55^, where maturation age (*Mat Age*) propensity of a genotype was evaluated using a logit link model, and a flexible threshold structure. We first selected the optimum model structure by evaluating a number of covariates using a semi-automated Akaike information criterion (AIC) selection approach in the MUMIN package in R^56^. In the *TAN* dataset a combination of four covariates (*SEX*, *FWS, FWG, SWG*) and their first order interactions were evaluated. In addition to these covariates, in the *NOR* dataset the geographical coordinates (i.e. latitude and longitude) were also evaluated in the model. For both datasets, the model with the lowest AIC value was chosen as the optimum model and applied in the GWA analysis (i.e. *FULL* model). *LEN* or *WGT* were not parametrised in the model, since these terminal traits are highly correlated to *mat age* (*Pearson’s r*, 0.89 and 0.88 for *LEN* and *WGT*, respectively. *P* < 10^-16^, for both) causing high co-linearity in the linear model, and also are not causatively linked to age at maturity. All covariates other than *SEX* were z-transformed to standard normal distribution, which prevented an unbalanced variance and covariance, and provided a better convergence in the maximum likelihood evaluation. In addition to analysing the *NOR* and *TAN* datasets separately, model selection was performed with the two datasets combined (*NOR+TAN,n* = 1,404) using parameters similar to those described above for *NOR* parametrization (i.e. geographical coordinates were included as parameters). Model selection was not performed for the Tornio samples, for which we did not have freshwater phenotypic information available and the GWA analysis was performed without phenotypic co-variates (i.e. *BASIC* model). Supplementary Table 2 lists the details of the model selection parameters for *TAN*, *NOR* and the combined (*TAN* + *NOR*) datasets.

All GWA analyses performed with the *FULL* model were also repeated with *BASIC* model to assess the effect of inclusion of covariates to the model (e.g.^57^, see Extended Data Fig. 1). The GWA analysis was performed using a model comparison approach, where the effect of the SNP loci was evaluated by comparing the likelihood of the observational level model (as above) to the additive genetic model with SNP loci as covariates. The genome-wide statistical significance was adjusted for multiple comparisons and genomic inflation (λ) for each analysis (*P = 0.05 × nSNP × λ*). Specific significance thresholds are listed in Extended Data Fig. 1.

To account population stratification, we fitted the same model as above but also included principle components (PC) derived from the genomic kinship matrix as fixed factors. Principle components were added sequentially in the model until origin of population no longer explained a significant portion of genetic variance across SNPs. Using principal components is a common and effective way to account for structured data when there are multiple populations in the data set^58^. Usually, principal components (PC) are added to the model sequentially until the genomic inflation (λ) factor drops below a certain threshold (e.g. λ< 1.05^58^). Sequential addition of PCs may be impractical with very large datasets composed of individuals from many populations when relatively complex models are used. Therefore, to assess the number of PCs to suitably account for population stratification, we quantified the reduction in among population variation (*σ*_*POP*_) upon subsequent addition of PCs. To do this analysis, within individual and population variation (*σ*_*ID*_, *σ*_*POP*_) were modelled as random terms in a generalized mixed model with a binomial link using the *glmer* function in the LME4 1.1-7 package^59^ in R, with or without including PCs as fixed effects. The minimum number of PCs in the model that the alterative hypothesis rejected (*H*_null_: *σ*_*POP*_ is significant) 95% of the time using 1,000 randomly selected SNPs was determined as the optimal number of PCs to account for population structure. The optimal number was one for *TAN*, and 14 for *NOR* and the combined (*TAN* + *NOR*) datasets (see Fig. 1 and Extended Data Fig. 1). Population structure in the *BAL* dataset was corrected using two PCs as fixed factors, which reduced the λ value to 1.07 (Extended Data Fig. 2). We also compared association statistics of BAL (*n* = 114) and the combined data (*TAN*+*NOR*, *n*=1,404) *post-hoc,* to assess the magnitude of the effect of sample size on the association statistic of the *VGLL3*_*TOP*_ locus. The *TAN*+*NOR* dataset was re-sampled 100,000 times with an equivalent sample size and age at maturity structure to the *BAL* dataset. The observed association statistic for the *VGLL3*_*TOP*_ locus in the BAL set was similar to that in the *TAN*+*NOR* re-sampled datasets (Kolmogorov-Smirnov test, *P* = 0.51, Extended Data Fig. 2) indicating that the lower P-value in *BAL* is likely due to the lower sample size.

### Identifying signatures of spatially divergent selection: F_ST_. outlier test

We used an extension of the Lewontin and Krakauer Test, the FLK test, which uses population trees (using Reynold’s genetic distances and neighbor joining algorithm) to estimate expected neutral evolution (null) among populations^60^. This method has been shown to perform well under different demographic scenarios^61^. The empirical null distribution of SNPs was identified using the estimated population tree and 100,000 simulations.

### Mode of inheritance and effect sizes of age at maturity loci

We detailed the genetic architecture of the age at maturity associated loci by evaluating the likelihood of several inheritance models. In addition to simple additive and dominance models, we also tested various models where dominance inheritance was modelled conditioned on sex. Extended Data Table 2 lists the details of each model. Models were compared using an information theocratic approach, where the model with the lowest AIC was accepted as the optimal model explaining the data. Coefficient of the optimal model for the *VGLL3*_*TOP*_ locus is given in Supplementary Table 3.

The patterns of dominance in the *VGLL3*_*TOP*_ locus were investigated in detail, at the unobserved liability scale. Deviations from additivity were tested for each sex separately. In addition, deviation of genetic architectures between the sexes was also tested. For these tests, we used genotype coefficients (β_*genotype*_) and the standard errors obtained by threshold model (Supplementary Table 3). We first standardized the coefficient to [0,1] range, such that (β_*LL*_ + β_*EE*_)/2 = 0.5 (i.e. the average of the homozygote genotypes). parametric permutations were drawn from genotype coefficients, and the additive expectation (i.e. null) was calculated as (β_*LL*_ + β_*EE*_)/2 (i.e. 100,000 parametric permutations). This was compared to β_*EL*_ (heterozygote). Test statistics were reported as the proportion of samples deviating from the null in one direction. For direct comparisons between sex dominance patterns, test statistics were reported as the proportion of samples deviating from the null (*H*_*null*_: β_*ELfemale*_ =β_*ELmale*_) in one direction.

### The proportion of variation explained by the VGLL3_TOP_ locus

To estimate the proportion of variance in age at maturity explained by the *VGLL3_TOP_* genotype, we employed an alternative modelling framework, where the response variable (*Mat Age*) is expressed at logit scale with the *y* = *Mat age*/(1 + *Mat Age*) transformation. Advantages of this transformation, where logit(*Mat Age* /(1 + *Mat Age*)) is equal to log(*Mat Age*) provides logistic coefficients at the (log) observational scale, are that it i) conveniently allows quantification of the examined variation in relation to total variation; and ii) enables quantification of effect size with a straightforward interpretation. We employed this framework to the *NOR*, *TAN*, and the combined datasets with the same specification as in the *FULL* model (plus including PCs as fixed factors) using *glmer* function in the LME4 package (Bates et al 2015) in R, using a binomial link and *bobyqa* optimizer. The model provided fits comparable to the additive cumulative proportional odds model, where, *R*^2^_*MF*_ and *R*^2^_*CS*_ were: 0.19 and 0.28 for the *TAN*, 0.13 and 0.17 for *NOR*, and 0.03, 0.12 for the combined dataset, respectively (See Supplementary methods and Supplementary Table 4 for model details).

Using this modelling framework, the proportion of variance explained by *VGLL3*_*TOP*_ genotype was estimated to be 42.8 % (s.e. 1.6), 37.5 % (s.e. 1.1), and 39.4 % (s.e. 1.1)for TAN, NOR and the combined datasets respectively, (1000 parametric bootstrapping, after accounting for populations structure), whereas sex specific estimates were, for females: 40.1 % (s.e. 2.2), 23.7 % (s.e. 1.2), and 30.5 % (s.e. 0.9), and for males: 40.1 % (s.e. 1.9), 32.2 % (s.e. 1.9), and 36.0 % (s.e. 1.8), for the *TAN*, *NOR* and the combined datasets, respectively. In addition, this modelling framework gave similar sex-specific dominance estimates to the additive cumulative proportional odds model (see Fig. 3a, Extended Data Figure 4, Supplementary Table 4).

In addition to sea age at maturity, we also analyzed the genotypic effect of *VGLL3*_*TOP*_ locus on variation in size at return, using ANOVA, and after accounting for maturation age. We modeled sexes separately due to the sex dependence genetic architecture. Population effects were calculated with a similar framework as in the GWA analysis such that the same number of PCs were used here to correct for the population inflation factor. Also, the genotype x maturity age interaction term was included into the models. In the combined dataset, 2.0 % (*P* = 0.0021) and 2.9 % (*P* = 4.80 × 10^-5^) of variation in length was explained by genotype at the *VGLL3*_*TOP*_ locus (Fig. 3b, Supplementary Table 1). Furthermore, there was a significant maturity age x genotype interaction in males with older age classes exhibiting higher length variation across genotypes (explaining 5.2 % of overall variation, *P* = 4.62 × 10^-7^, Fig. 3b, Supplementary Table 1). Supplementary Table 1 summarizes the model parameters, for the *TAN, NOR* and the combined datasets.

Genome resequencing and functional variant detection. Thirty-two wild Atlantic salmon were selected for whole genome resequencing from seven populations (three from the Barents/White Sea and four from the Atlantic phylogeographical group, see also Extended Data Table 1, and Supplementary Table 5). Three individuals per population were resequenced, except for the Tana sub-populations, where 14 individuals were resequenced. DNA was isolated from 14 adipose fin-clips (stored in ethanol) and 18 scale samples collected in 2012 and 2013 (stored in paper envelopes) using Qiagen DNAeasy (Qiagen, Germany) kits according to manufacturer’s recommendations. DNA was quantified using Qubit fluorometry (Invitrogen, USA).

For high-quality adipose tissue derived DNA and two scale derived DNA extractions that had high DNA quantity and quality, sequencing libraries were produced using the TruSeq DNA PCR-free Library Preparation Kit. Libraries for the remaining 16 scale derived DNA extracts were prepared using the TruSeq Nano DNA Library Preparation Kit. The main motivation for this was to select kits most suited to available sample quantities, both kits use mechanical fragmentation (Covaris) thus limiting a bias caused by using a mixture of enzymatic and mechanical approaches. Library preparations were performed according to manufacturer’s instructions (Supplementary Table 5). All libraries were subjected to a fragment size selection (mode = 350bp) and sequenced to generate 2 × 125nt paired-end reads using an Illumina HiSeq 2500 platform. Sample preparation and sequencing was performed by the Norwegian Sequencing Centre, Ullevål (Oslo, Norway). Only reads passing Illumina’s chastity filter were used in subsequent analysis. We further used FastQC to assess sequencing quality, passing lanes where the per-base quality score box plot indicated bases 1-110 having > Q20 for > 75 % of the reads. All lanes passed the quality criteria.

Reads were mapped to the salmon reference genome (NCBI WGS accession number AGKD04000000) using BWA mem version 0.7.10-r789^62^. The thirty-two samples were sequenced to a depth of around 18X (8-32X). In total, 7.6 billion out of 8.3 billion reads (92 %) were properly aligned to the genome. SNPs and short indels were identified using Freebayes version v0.9.15-1^63^). To filter away low-quality variants, we used the run-time parameters – *use-mapping-quality* and –*min-mapping-quality 1*, in addition to ‘*vcffilter* -f “QUAL > 20”’. SNPs and short indels were annotated using snpEff version 4.0e^64^. The snpEff annotation database was based on the CIGENE annotation v2.0 (manuscript in prep.).

### Selection analysis

Genotypic data from all individuals (*n* = 1,518) were phased using the Beagle 4.0 software^65^ with imputation for missing genotypes (% 0.2 of calls) using the parameter window size = 50,000 and overlap size = 3,000. 10, 40 and 50 iterations were parameterized for burn-in, phasing and imputation of the data, respectively, and physical distances were used as a proxy for genetic distances. We used an extended homozygosity haplotype (*EHH)*^66^ based test to detect footprints of selection, using the REHH v3.1.1 package ^67^. We first computed integrated *EHH* scores (*iHH*) using the *scan_ehh* function in REHH v3.1.3 with default parameters. We then computed the integrated haplotype scores per population (*iHS*)^68^ using the *ihh2ihs* function in REHH v3.1.3 (frequency bin = 0.05, *maf* = 0.05). *iHS* is a metric to quantify the difference in extended haplotype homozygosity (EHH) at a given SNP between the two alleles. The *iHS* score is standardized empirically to the distribution of observed *iHS* scores over a range of SNPs with similar derived allele frequencies. Throughout the haplotype analyses we did not define the evolutionary state of the alleles (i.e. ancestral versus derived) as it was not of relevance for this study. Therefore the sign of the *iHS* score is arbitrary at every SNP, where the higher frequency allele in the initial phasing stage is coded as the ancestral allele.

We tested whether variation in age at maturity among populations could be maintained by selection towards an optimum age at maturity composition within each population given gene flow is expected among the phenotypically divergent local populations sampled. In order to test for divergent local selection, we employed a linear model where the median *iHS* scores were regressed over the average sea age of the populations (Extended Data Table 1). Statistical significance was assessed by comparing the regression coefficient (i.e. the proportion of variation in the *iHS* score explained by sea age) at the locus of interest to the null distribution at the genome wide level. For this analysis, we calculated the *iHS* statistics for every population with at least 16 successfully genotyped individuals (32 haplotypes), and used an equal number of individuals per population (by random selection of individuals if *n* > 16). We assessed the effect of using 16 randomly selected individuals from the two populations in the *TAN* dataset and *iHS* scores in the reduced set were in good agreement with the full dataset (*Pearson’s r = 0.72* and *0.75* for younger and older age structured sub-populations, respectively. *P* < 10^-16^ for both datasets), suggesting the robustness of *iHS* analysis with the applied sample size (Extended Data Fig. 5e,f).

